# Early Inhibition of Retinoic Acid Signaling Rapidly Generates Cardiomyocytes Expressing Ventricular Markers from Human Induced Pluripotent Stem Cells

**DOI:** 10.1101/856575

**Authors:** Pranav Machiraju, Joshua Huang, Fatima Iqbal, Yiping Liu, Xuemei Wang, Chad Bousman, Steven C. Greenway

**Affiliations:** Department of Pediatrics, Cumming School of Medicine, University of Calgary, Calgary, AB T2N 4N1, Canada; Centre for Advanced Technologies, Cumming School of Medicine, University of Calgary, Calgary, AB T2N 4N1, Canada; Hotchkiss Brain Institute, Cumming School of Medicine, University of Calgary, Calgary, AB T2N 4N1, Canada; Department of Medical Genetics, Cumming School of Medicine, University of Calgary, Calgary, AB T2N 4N1, Canada; Department of Psychiatry, Cumming School of Medicine, University of Calgary, Calgary, AB T2N 4N1, Canada; Department of Physiology and Pharmacology, Cumming School of Medicine, University of Calgary, Calgary, AB T2N 4N1, Canada; Department of Biochemistry & Molecular Biology, Cumming School of Medicine, University of Calgary, Calgary, AB T2N 4N1, Canada; Alberta Children’s Hospital Research Institute, Cumming School of Medicine, University of Calgary, Calgary, AB T2N 4N1, Canada; Department of Cardiac Sciences, Cumming School of Medicine, University of Calgary, Calgary, AB T2N 4N1, Canada; Libin Cardiovascular Institute of Alberta, Cumming School of Medicine, University’ of Calgary, Calgary, AB T2N 4N1, Canada

## Abstract

**SUMMARY:** Current protocols for the differentiation of cardiomyocytes from human induced pluripotent stem cells (iPSCs) generally require prolonged time in culture and result in heterogeneous cellular populations. We present a method for the generation of beating cardiomyocytes expressing specific ventricular markers after just 14 days. Addition of the pan-retinoic acid receptor inverse agonist BMS 493 to human iPSCs for the first 8 days of differentiation resulted in increased protein expression of the ventricular isoform of myosin regulatory light chain (MLC2V) from 18.7% ± 1.72% to 55.8% ± 11.4% (p <0.0001) in cells co-expressing the cardiac muscle protein troponin T (TNNT2). Increased MLC2V expression was also accompanied by a slower beating rate (49.4 ± 1.53 vs. 93.0 ± 2.81 beats per minute, p <0.0001) and increased contraction amplitude (201% ± 8.33% vs. 100% ± 10.85%, p <0.0001) compared to untreated cells. Improved directed differentiation will improve *in vitro* cardiac modeling.

## INTRODUCTION

Reprogramming of human somatic cells generates patient- and disease-specific induced pluripotent stem cells (iPSCs) that can be subsequently differentiated into tissue-specific cells to enable *in vitro* modelling of common and rare human diseases (Takahashi et al., 2007). Protocols to differentiate iPSCs into cardiomyocytes (iPSC-CMs) have become highly efficient with commercially available kits now able to generate cells that almost universally (>90%) express the cardiac muscle-specific protein troponin T (TNNT2) (Karakikes et al., 2015; Lian et al., 2013). However, these iPSC-CMs are still relatively heterogeneous since they contain multiple cardiomyocyte subtypes, including atrial, ventricular, and pacemaker cells (Lee et al., 2017). This heterogeneity can confound disease modelling and potentially limits future clinical application.

Recently, efforts have been made to improve the specification of iPSC-CM differentiation protocols to result in the generation of specific cardiac subtypes (Lee et al., 2017; Pei et al., 2017). The retinoic acid receptor (RAR) pathway has emerged as a key regulator in cardiomyocyte differentiation with retinoic acid signalling specifying the development of atrial myocytes and antagonism of the RAR pathway favouring the development of ventricular myocytes (Devalla et al., 2015; Lee et al., 2017; Zhang et al., 2011).

We have developed a simple and reproducible methodology to improve the generation of cardiomyocytes demonstrating ventricular-specific properties using the pan-RAR inverse agonist BMS 493 and a commercially available cardiomyocyte kit from STEMCELL Technologies. Furthermore, these ventricular myocytes can be generated relatively quickly, requiring just 14 days in culture following the initiation of differentiation.

## RESULTS

We used three distinct human iPSC lines (Gibco, CIM001 and CIM008) for all experiments. All lines were obtained from individuals without known or active cardiac disease. All three lines were differentiated into beating iPSC-CMs using the Cardiomyocyte Differentiation Kit (Catalog #05010) from STEMCELL Technologies (Vancouver, Canada). Single iPSCs seeded onto Matrigel and maintained with mTeSR Plus medium became >98% confluent after 2 days in culture. Cells were then either treated with vehicle control, BMS 493 (1 μM) from days 4-8 of differentiation or BMS 493 (1 μM) from days 0-8 of differentiation (Figure S1).

To quantify the generation of ventricular cardiomyocytes, flow cytometry was used to determine the number of cells expressing TNNT2 and the ventricular isoform of myosin regulatory light chain (MLC2V), specific for human ventricular cells (Franco et al., 1998; Franco et al., 1999) after 14 days in culture following differentiation in the presence and absence of BMS 493 (Figures 1 and S2). Nearly all (94.5% ± 1.10%) untreated iPSC-CMs expressed TNNT2 as did cells treated with BMS 493 from days 4-8 (95.2% ± 0.67%). Interestingly, all three iPSC lines differentiated in the presence of BMS 493 from days 0-8 demonstrated significant reductions in TNNT2 expression, ranging from 63.5% ± 3.42% for CIM001 to 83.1% ± 2.86% (Gibco) and 82.6% ± 4.23% for CIM008 (Figure 1G). However, those cells incubated with BMS 493 from days 0-8 of differentiation did demonstrate increased generation of ventricular-specific cardiomyocytes with 44.4% ± 4.64%, 61.5% ± 6.97%, and 65.3% ± 7.22% of cardiomyocytes expressing MLC2V for the Gibco, CIM001 and CIM008 iPSC-CMs, respectively, compared to only 18.7% ± 1.72% of untreated cells and only 17.3% ± 1.39% of iPSCs treated with BMS 493 from days 4-8 of differentiation (mean ± SD, Figure 1H).

**Figure 1.**
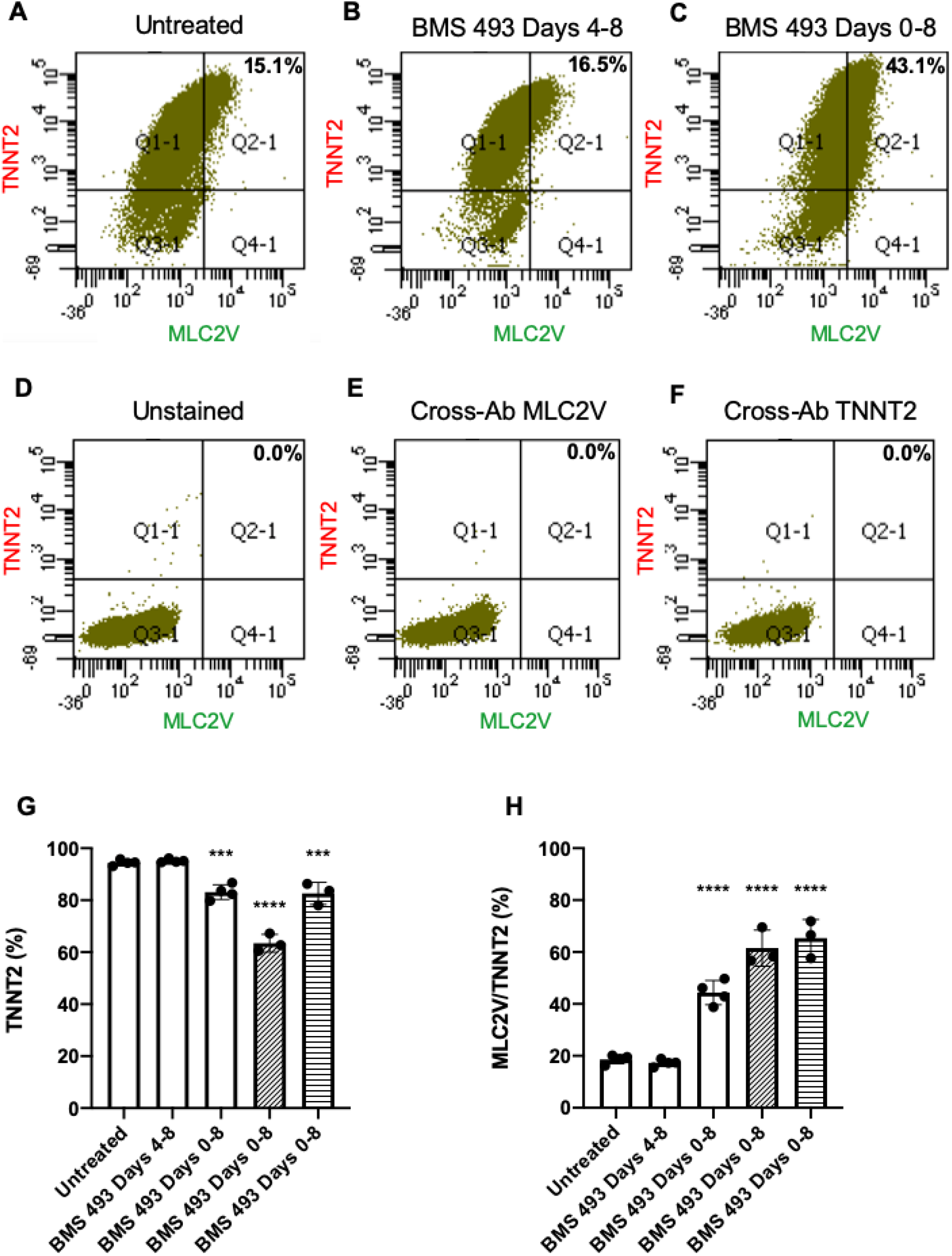
Increased generation of iPSC-CMs expressing MLC2V after incubation with BMS 493 from days 0-8 of differentiation. (A-C) Representative flow cytometry quadrant graphs showing the relative proportion of cells expressing TNNT2 and MLC2V without (15.1%) and with BMS 493 for days 4-8 (16.5%) or days 0-8 (43.1%) of differentiation for the Gibco line of iPSCs. (D-F) Flow cytometry quadrant graphs for unstained or cross-antibody (mismatch of secondary antibody to primary antibody) controls for Gibco iPSCs. Gates for all graphs are based on unstained and cross-antibody controls. Data represent sorting of 50,000 cells per group. Percentages of treated and untreated iPSC-CMs expressing TNNT2 after 14 days in culture. Percentages of treated and untreated iPSC-CMs positive for TNNT2 expressing MLC2V after 14 days in culture. Data are mean ± SD from n = 3-4 independent experiments using Gibco (white bars), CIM001 (bars with hatched lines) and CIM008 (bars with horizontal lines) iPSC-CMs. ***p <0.001, ****p <0.0001.

Cell morphology changed drastically during the initial cardiomyocyte differentiation phase (days 0-8) with cells displaying a cardiac-like morphology (long and large cells cross-linked to each other) being visible from day 8 onwards. Three-dimensional cardiac structures became visible after day 10 and remained for the duration of culture in all cell groups due to the presence of a Matrigel sandwich (Zhang et al., 2012). Untreated cardiomyocytes and cardiomyocytes treated with BMS 493 from days 4-8 of differentiation started beating around day 8 of differentiation with the amount of contractile activity increasing between days 10-14. In contrast, cells treated with BMS 493 from days 0-8 started contracting only after days 10-12 of culture. However, contractility in these cells appeared to be slower and more forceful compared to the other two treatment groups with contractile centers also appearing larger.

To assess the contractile phenotype of the iPSC-CMs, beating rate and contraction amplitude were measured from videos of contracting cardiomyocytes visualized using phase-contrast microscopy. The mean beating rate (beats per minute) of iPSCs treated with BMS 493 from days 0-8 of differentiation was significantly lower than untreated cells. The average beating rate (± SEM) for the Gibco, CIM001 and CIM008 cell lines were 36.5 ± 2.33, 52.74 ± 0.43, and 58.5 ± 1.48 bpm, respectively, compared to 93.0 ± 2.81 and 85.0 ± 4.78 for untreated cells and cells treated with BMS 493 from days 4-8 (Figure 2A). Cardiomyocytes from all three cell lines treated with BMS 493 from days 0-8 showed significantly higher contraction amplitudes compared to untreated cells or iPSCs differentiated in the presence of BMS 493 from days 4-8 (Figure 2B). Gibco, CIM001 and CIM008 cell lines treated from days 0-8 with BMS 493 showed 217.7% ± 18.80%, 197.3% ± 24.89%, and 189.8% ± 14.46% (mean of untreated ± SEM) respectively (Figure 2B).

**Figure 2.**
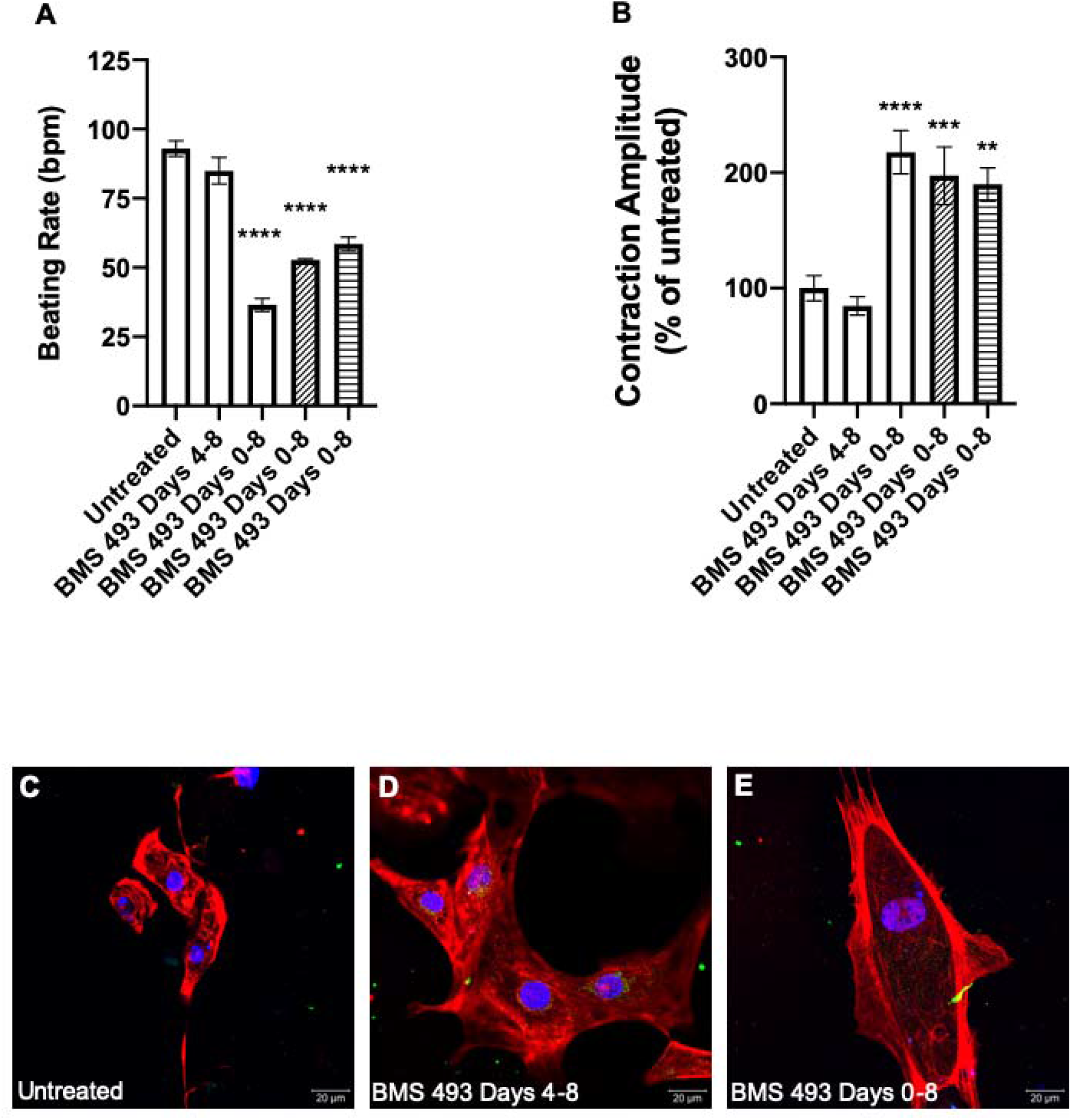
Decreased beating rate, increased contraction amplitude of iPSC-CMs after treatment with BMS 493 during days 0-8 of differentiation. (A) Average beating rate (beats per minute, bpm) for iPSC-CMs with and without BMS 493 treatment. (B) Contraction amplitude of iPSC-CMs from each treatment group relative to untreated cells. Data are mean ± SEM for n = 25-29 cardiomyocyte colonies using Gibco (white bars), CIM001 (bars with hatched lines) and CIM008 (bars with horizontal lines) iPSC-CMs. Cell properties were measured after a total of 14 days in culture. **p <0.01, ***p <0.001, ****p <0.0001. (C-E) Immunocytochemistry of single-cell iPSC-CMs (untreated, treated with BMS 493 from days 4-8 or treated with BMS 493 from days 0-8 of differentiation) stained for TNNT2 (red), MLC2V (green) and DAPI (blue). Scale bar indicates 20 μm.

Immunocytochemistry of untreated cells and cells treated with BMS 493 from days 4-8 or 0-8 of differentiation was performed after a total of 14 days in culture. Single-cell images show high levels of TNNT2 protein in all three treatment groups. Qualitatively, there is also an increase in intracellular MLC2V protein in cardiomyocytes treated with BMS 493 from days 0-8, consistent with our flow cytometry data. Additionally, cardiomyocytes treated with BMS 493 from days 0-8 appear to be larger in size and more elongated than the other cardiomyocytes (Figures 2C-E).

## DISCUSSION

In this study, we describe the rapid and improved generation of ventricular-specific cardiomyocytes from human iPSCs using a commercially available kit in the presence of BMS 493 from days 0-8 of differentiation.

Previous work has described the importance of RAR signalling (Collop et al., 2006; Pan and Baker, 2007) and even identified the potential utility of BMS 493 in cardiomyocyte generation (Pei et al., 2017). However, the study by Pei et al. was heavily time-dependent with their cultured iPSC-CMs only showing the increased levels of MLC2V protein expression approaching those seen in our study after >60 days in culture (Pei et al., 2017). Furthermore, their differentiation protocol may lack reproducibility in comparison to a purchased kit.

The timing of BMS 493 exposure during differentiation appears to be critical, with exposure during the first 4 days required for increased MLC2V expression. Interestingly, BMS 493 exposure from days 0-8 was associated with relatively decreased TNNT2 expression. In the future, this reduction could potentially be avoided through the addition of a lactate-purification step to eliminate any non-cardiac cells in culture (Tohyama et al., 2013). However, as ascertained by rigorous flow cytometric testing, all three cell lines treated with BMS 493 from days 0-8 of differentiation showed a significantly higher proportion of cardiomyocytes expressing MLC2V. Our flow cytometry data was supported through immunocytochemistry staining of the iPSC-CMs. Based on previous literature, we hypothesize that other non-ventricular cardiomyocytes are likely atrial or pacemaker cells (Lee et al., 2017; Pei et al., 2017).

A key hallmark of ventricular cardiomyocytes is their slower beating rate and higher contractile force in comparison to other cardiomyocyte sub-types. Our data, ascertained from multiple cardiomyocyte colonies in each treatment group, suggests that our protocol resulted in a higher proportion of cells with contractile function comparable to ventricular cardiomyocytes *in vivo.* The higher contraction amplitude and lower beating rate observed in our treated cells may also indicate a change in the maturation state towards more adult-like cardiomyocytes (Denning et al., 2016; Karakikes et al., 2015). However, although the specification and maturation of cardiomyocytes may be interconnected, they are not mutually inclusive. Currently, prolonged time in culture appears to be the predominant factor responsible for converting fetal-like iPSC-CMs into cells with the characteristics of adult cardiomyocytes (Karakikes et al., 2015; Machiraju and Greenway, 2019). Further studies into the maturation state of our cells will need to be undertaken. It is likely that without the implementation of specific maturation approaches (e.g. mechanical or metabolic manipulation) they cannot be defined as mature ventricular cardiomyocytes.

In summary, combining the current iteration of the cardiomyocyte differentiation kit from STEMCELL Technologies and the pan-RAR inverse agonist BMS 493 from days 0-8 of differentiation, we are able to increase the proportion of iPSC-CMs expressing ventricular-specific markers within a relatively short culture time of 14 days. The combination of our protocol with additional strategies to promote specification and/or maturation will likely yield further positive results.

## ACKNOWLEDGEMENTS

This work was supported by the Alberta Children’s Hospital Research Institute, the University of Calgary and the Children’s Cardiomyopathy Foundation (to S.C.G.).

## AUTHOR CONTRIBUTIONS

P.M. and S.C.G. designed the experiments and wrote the manuscript. P.M., J.H., F.I., Y.L. and X.W. performed the experiments. C.B. provided essential reagents. P.M. performed the data analysis. S.C.G. supervised the project and provided funding.

---

**Supplemental Figure 1.**
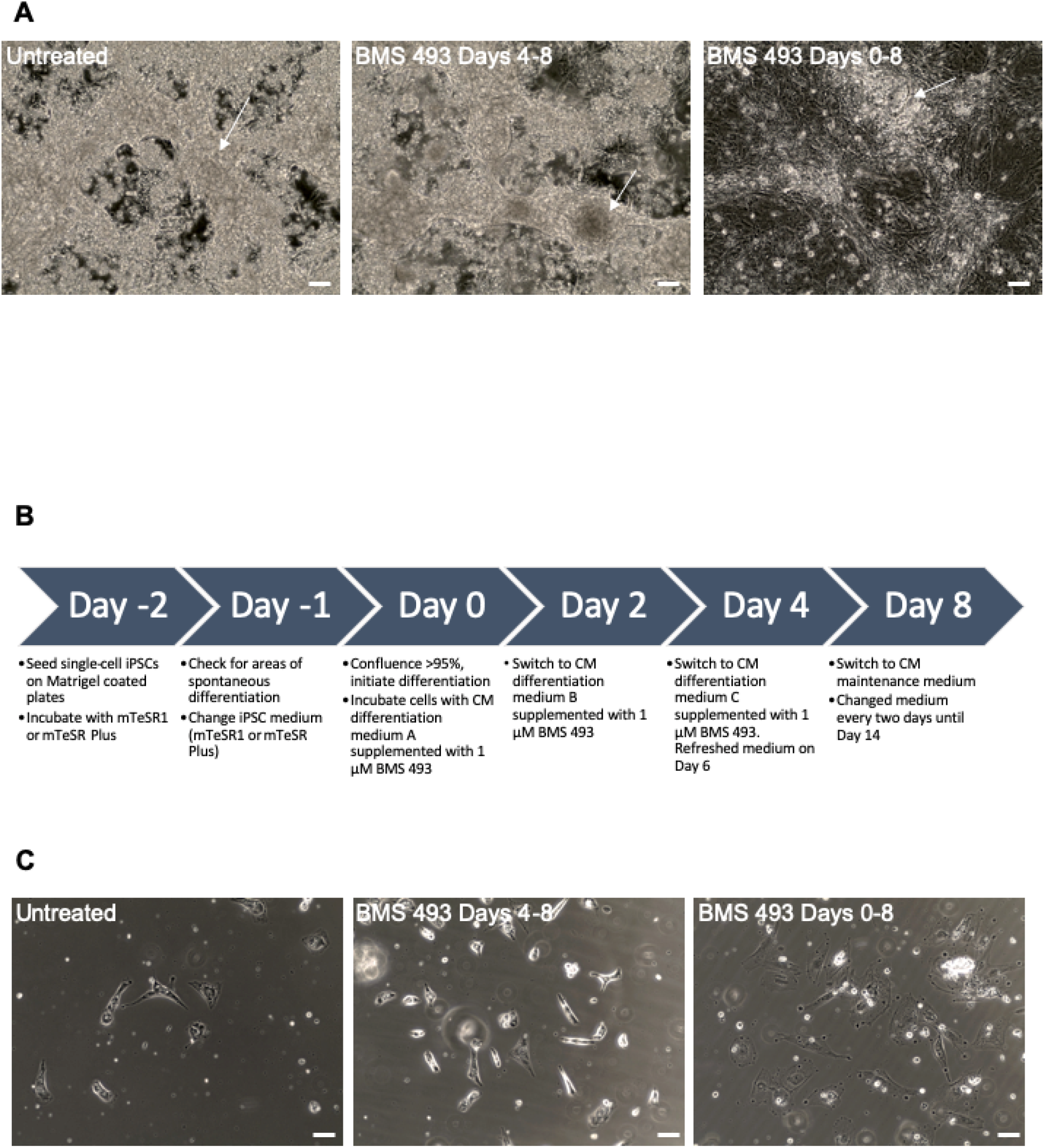
Cells and experimental protocol. (A) Phase-contrast images of iPSC-CMs (untreated, treated with BMS 493 from days 4-8 and treated with BMS 493 from days 0-8 of differentiation). Arrows indicate spontaneously contracting cardiomyocyte centers. Scale bar indicates 80 μm. (B) Flow chart outlining the differentiation protocol used in this study. Day 0 marks the beginning of cardiomyocyte differentiation followed by serial media changes. (C) Phase-contrast images of single-cell hPSC-CMs (untreated, treated with BMS 493 from days 4-8 and treated with BMS 493 from days 0-8 of differentiation). Scale bar indicates 40 μm.

**Supplemental Figure 2.**
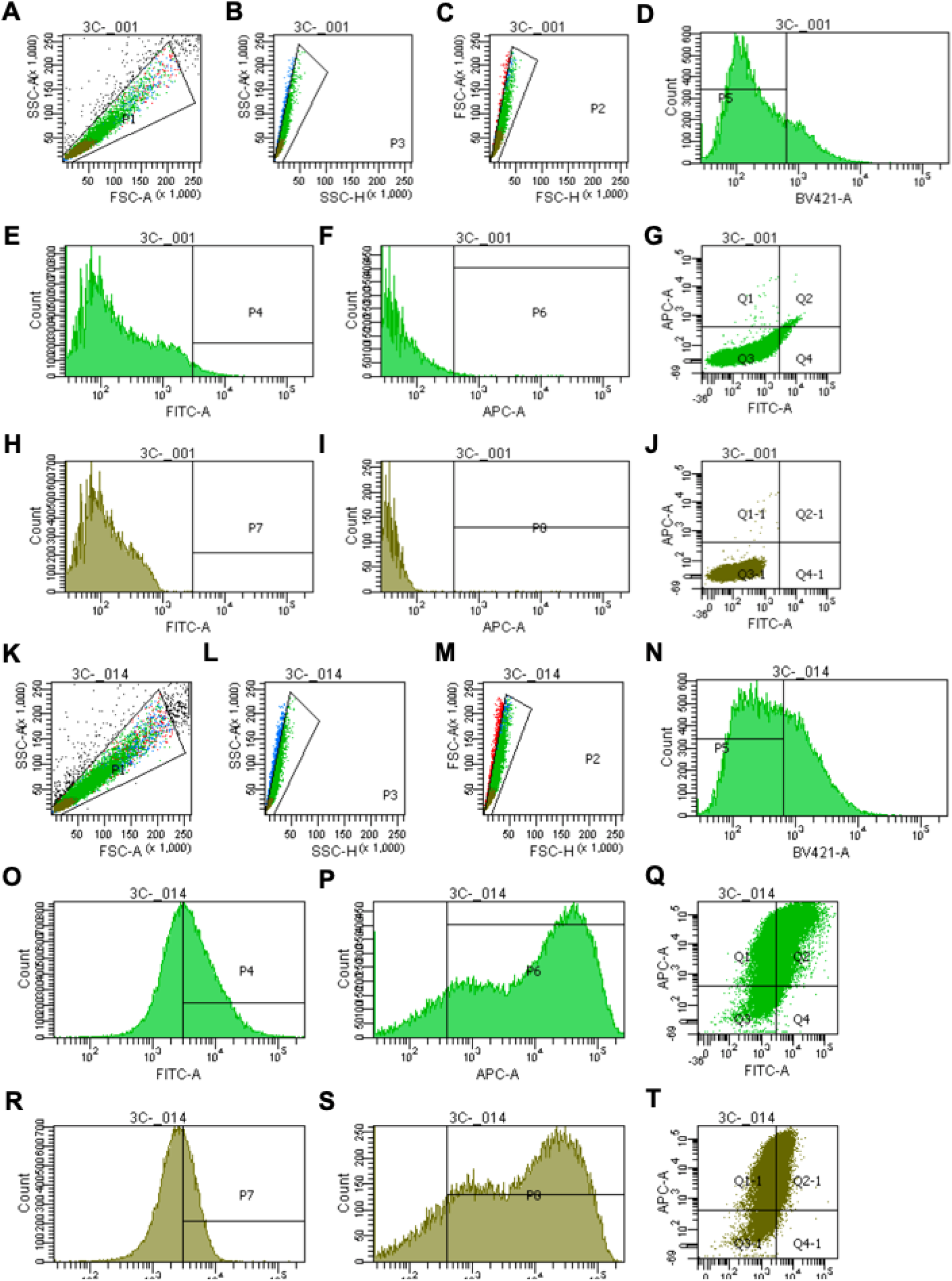
Flow cytometry gating strategy. (A-C) Series of gates showing doublet-discrimination gating for unstained control cells. (D) An empty laser channel (bright violet) was used to separate dead cells and non-specific fluorescence. (E-G) Histograms and quadrant graph of unstained cells after doublet-discrimination gating. (H-J) Histograms and quadrant graph of unstained cells after doublet-discrimination gating and removal of dead cells through the bright violet empty channel. (K-T) Flow cytometry graphs indicating the gating strategy used on one stained positive replicate. (K-M) Series of gates showing doublet-discrimination gating. (N) Bright violet empty channel to sort out dead cells and non-specific fluorescence. (O-Q) Histograms and quadrant graph of stained cells after doublet-discrimination gating. (R-T) Histograms and quadrant graph of stained cells after doublet-discrimination gating and removal of dead cells through the bright violet empty channel. Data represent sorting results for 50,000 cells. FITC and APC lasers were used to analyze MLC2V and TNNT2, respectively.

## METHODS

### LEAD CONTACT AND MATERIALS AVAILABILITY

Further information and requests for resources and reagents should be directed to and will be fulfilled by the Lead Contact, Steven Greenway. This study did not generate new or unique reagents.

### EXPERIMENTAL MODEL AND SUBJECT DETAILS

Three human iPSC lines were used in this project. The first line, Gibco, was an Episomal iPSC line purchased from Thermo Fisher Scientific (cat. A18945). The other two lines (CIM001 and CIM008) were iPSCs reprogrammed from peripheral blood mononuclear cells using Sendai virus at the University of Melbourne core facility. All iPSC lines were cultured for 10 passages using standard iPSC protocols prior to differentiation.

## METHOD DETAILS

### Culture and differentiation of iPSCs into cardiomyocytes (iPSC-CMs)

All iPSCs were grown on Matrigel with mTeSR 1 or mTeSR Plus media to preserve pluripotency. Clump passaging was performed using EDTA (Thermo Fisher Scientific, cat. 15575020) while single-cell passaging was accomplished using Accutase (Thermo Fisher Scientific, cat. A1110501). Prior to cardiomyocyte differentiation, iPSCs were seeded as single cells in mTeSR Plus medium containing a 1:100 dilution of RevitaCell (Thermo Fisher Scientific, cat. A2644501) to limit cell death. A split ratio of 1:2.5 was used to achieve over 95% confluency two days post-seeding onto Matrigel-covered cell-culture plates. Following a visual check to ensure proper iPSC morphology, cardiomyocyte differentiation was initiated. To differentiate iPSCs into cardiomyocytes, the Cardiomyocyte Differentiation Kit (cat. 05010) from STEMCELL Technologies (Vancouver, Canada) was used. The kit protocol was followed exactly as outlined by the manufacturer. On Day 0, confluent iPSCs were incubated with 5 mL of complete medium A plus a 1:100 dilution of Matrigel, for 48 hours. On Day 2, medium was changed to 5 mL of complete medium B for an additional 2 days. On Days 4 and 6, 5 mL of complete medium C was added to all wells and left to incubate for 48 hours. Cardiomyocytes were cultured in 6-well plate wells resulting with a medium volume of 5 mL. BMS 493 (TOCRIS, cat. 3509), a pan-RAR inverse agonist, was either not used, used from days 4-8 of differentiation, or used from days 0-8 of differentiation at a final concentration of 1 μM. After day 8, differentiation was complete and cells were cultured in complete maintenance medium (changed every two days) until Day 14. At Day 14, cardiomyocytes were isolated for all experiments. All cells were cultured in a sterile, mycoplasma-free environment without the use of antibiotics. Gibco iPSCs were used to evaluate the time-course of BMS 493 supplementation and iPSC lines CIM001 and CIM008 were only differentiated with the addition of BMS 493 from days 0-8 of differentiation to validate and replicate our findings.

### Flow cytometry

Expression of cardiac- and ventricular-specific markers were quantified through flow cytometry. Briefly, cells were dissociated using the Cardiomyocyte Dissociation Kit (STEMCELL Technologies). Next, 7-8 mL of 2% paraformaldehyde (PFA, J.T. Baker) was added and the solution was allowed to fix at room temperature (RT) for 15 min. After fixation, the cell suspension was centrifuged at 300 × *g* for 9 min at RT. The supernatant was then aspirated and 7-8 mL of cold 90% methanol was added. The mixture was incubated at 4°C on ice for 15 min. Following incubation, the solution was centrifuged again at 300 × *g* for 9 min. After two washes with 0.25% BSA (flow buffer), cells were centrifuged at 300 × *g* for 9 min. Flow buffer was added and cells were distributed into 2 mL microcentrifuge tubes. Tubes were centrifuged at 200 × *g* for 6 min at RT. The supernatant was aspirated, and the cells were re-suspended in 1.5 mL of flow buffer. Cells were then centrifuged and 1 mL of 0.1% Triton X-100 was added for 15 min to permeabilize the cells. After one more centrifugation step and aspiration, 1 mL of 10% BSA was added as a blocking agent for 20 min at RT. Next, after centrifugation and aspiration, 0.2 mL of flow buffer containing 1:400 TNNT2 (R&D Systems, cat. MAB1874) and 1:100 MLC2V (ProteinTech, cat. 10906-1-AP) antibodies was added to each positive replicate tube. The suspension was then incubated for 1 hour at RT. Following incubation, the tubes were centrifuged at 200 × *g* for 5 min at RT. The supernatant was then aspirated and 1 mL of 10% BSA was added to prevent non-specific binding of the primary antibodies. Next, after centrifugation and aspiration, cells were washed in flow buffer three times with cells being incubated in flow buffer for 5 min for each wash. The suspension was then centrifuged and 0.2 mL of flow buffer containing the secondary antibodies (ThermoFisher, AlexaFluor 488, cat. A21121 and AlexaFluor 647, cat. 32728) was added at a concentration of 1:2000 for each secondary antibody. The samples were incubated for 30 min at RT while protected from light. Following incubation, the tubes went through a series of three wash steps with 1.5 mL flow buffer followed by centrifugation at 200 × *g* for 5 min each time. Once ready to analyze, cells were centrifuged, the supernatant was aspirated and 300 μL of flow buffer was added per tube. The solutions were then transferred to clear flow cytometry tubes and the solution was agitated to ensure a homogenous cell suspension.

Flow cytometry analysis was performed in the Flow Cytometry Core Facility at the University of Calgary. The experiments used a variety of controls including unstained, cross-antibody control (each primary antibody with mismatched secondary antibody) and secondary antibody-only controls. Doublet discrimination gating was performed, and gates were assigned based on the cross-antibody controls after exclusion of dead cells through an empty channel (bright violet). A total of 50,000 cells for each replicate (n = 4 for Gibco iPSC-CMs and n = 3 for CIM001 and CIM008 iPSC-CMs) was quantified for each treatment condition. Quadrant graphs were generated in BD FACS Diva software. Bar graph data are presented as mean ± SD. Each cell line was analyzed based on its own controls to ensure accuracy.

### Immunocytochemistry

To prepare cells for immunocytochemistry, cardiomyocytes were isolated at day 14 post-induction of cardiomyocyte differentiation. A total of 40,000 cardiomyocytes were seeded onto individual, sterilized microscope coverslips placed on the bottom of a 24-well tissue culture plate. Cells were then allowed to grow for 24 hours prior to fixation. Cells on glass coverslips were washed twice with Dulbecco’s phosphate-buffered saline (DPBS) then fixed with 4% PFA in DPBS and pre-incubated at 37°C for 15 min. Cells were then washed three times with DPBS, quenched with 50 mM NH_4_Cl for 15 min at RT then washed again with DPBS and stored at 4°C. When ready to stain, cells were permeabilized with 0.1% Triton X-100 (Sigma-Aldrich, cat. 9002-93-1) in PBS for 15 min then washed three times with DPBS, blocked with 10% FBS for 25 min at RT then incubated with 1:50 TNNT2 and 1:100 MLC2V primary antibodies diluted in 5% FBS for 1 hour at 37°C. Cells were washed three times (five minutes per wash) with 5% FBS diluted in DPBS. Cells were then incubated with the AlexaFluor 488 and AlexaFluor 647 secondary antibodies (1:1000, Thermo Fisher Scientific) in 5% FBS for 1 hour at RT. Cells were washed then stained with NucBlue Fixed Cell Reagent (Thermo Fisher Scientific, cat. R37606) for 5 minutes then washed. Next, coverslips were placed onto glass microscope slides using Prolong Gold Anti-fade (Thermo Fisher Scientific, cat. P36930) and left overnight at RT before being imaged on a Zeiss LSM880 confocal microscope using a 63× oil objective. All treatment conditions were imaged using the same microscope settings including laser intensity, field of view, and magnification.

### Quantification of contractile characteristics

The contraction amplitude of beating iPSC-CMs was calculated using ImageJ and the MUSCLEMOTION plugin (Sala et al., 2018). Videos of iPSC-CMs were taken using an iPhone at 1080p, 60 frames per second and a phase-contrast microscope at 10× magnification. The videos were then split into image sequences using the MPEGStreamclip software. To optimize settings for each image sequence analyzed, a reference frame from each contractile colony was selected. The automatic reference frame was set manually along with the beat width. The software was then run for 25 beating colonies per treatment condition and contraction amplitude was recorded and plotted in Prism 8 (GraphPad Software). Data are presented as mean ± SEM with n = 26 except for CIM001 and CIM008 which are n = 26 and 29, respectively.

The beating rate of the iPSC-CMs was calculated manually. The number of beats in each video of a contracting colony over 15 seconds was recorded. This number was multiplied by four in order to ascertain beats per minute (bpm). A total of 25 colonies per treatment condition were analyzed. Data are presented as mean ± SEM with n = 25 except for CIM001 and CIM008 which are n = 27 and 26, respectively.

### QUANTIFICATION AND STATISTICAL ANALYSIS

Data are presented as means with standard deviation (SD) or standard error (SEM). Data were plotted and analyzed using GraphPad Prism using one-way ANOVA and unpaired t-tests as appropriate where n signifies the number of cells used or the number of independent experiments.

